## Supplementary Figures and Tables for "An aphid-resistant plant metabolite as a candidate aphicide: Insight into the bioactivity and action mode of betulin against aphids"

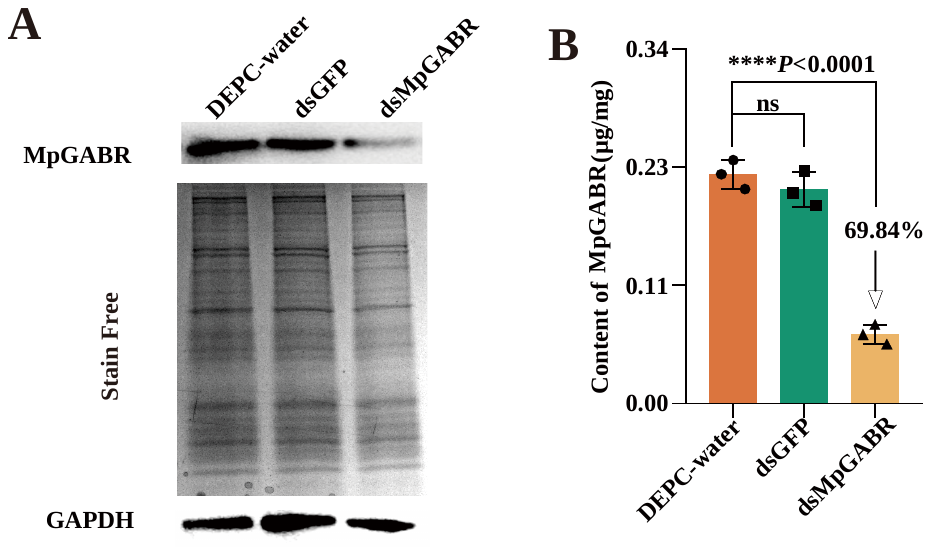

**Figure S1.** Protein expression of MpGABR after RNAi. Western blotting analysis (A) of MpGABR expression after RNAi at 48 h post-dsRNA feeding, relative (B) to expression level after DEPC-water treatment.

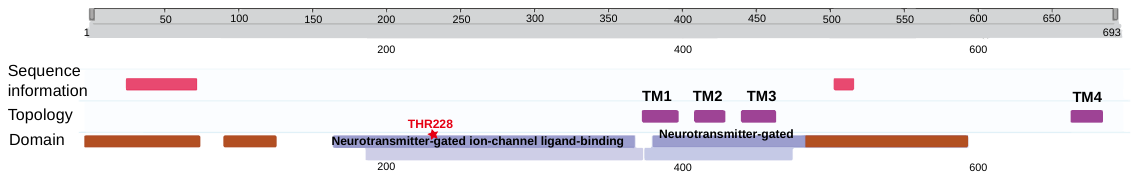

**Figure S2.** Betulin binding site on the domain of MpGABR. The structure of MpGABR protein was predicted on UniProt website. Pink bars indicate basic residue sequences and brown bars mean disordered regions.

**Table S1.** LC_50_ values of betulin and pymetrozine against *M. persicae* at 48 h

| **Compounds** | **Regression equation** | **LC_50_**  **(mg⋅mL^−1^)** | **95% Confidence**  **Interval (mg⋅mL^−1^)** | ***r^2^*** |
| --- | --- | --- | --- | --- |
| betulin | Y=6.9323+2.4621X | 0.1641 | 0.1635-0.1648 | 0.9235 |
| pymetrozine | Y=4.9840+0.6195X | 1.0612 | 1.0037-1.1221 | 0.9806 |

**Table S****2.** Primers used for cloning, RNAi, heterologous expression, and qRT-PCR

| **Experiment** | **Gene ID** | **Gene** | **Primer names and sequences (5′-3′)** | **Product length (bp)** | |
| --- | --- | --- | --- | --- | --- |
| qPCR | 111036118 | MpGABR (GABAA receptor) | F: TCGTCAACGAGAAACAGTCG | | 229 |
|  |  |  | R: GAGTTTGGTCCCTTGTTCCA | |  |
|  | 111041856 | MpGABRAP (GABAA receptor-associated protein) | F: CTGGTACCCTCCGACCTG | | 202 |
|  |  |  | R: GTCCATAGACATTTTCAT | |  |
|  | 111036117 | MpGABRB (GABAA receptor subunit beta) | F: TGAAGTCGGTGTCACCATGT | | 232 |
|  |  |  | R: GGTGGTCGCTATGTGGAAGT | |  |
|  | 111028237 | maltase 2-like | F: TTTCGGCAGTGAAACTGAAA | | 164 |
|  |  |  | R: CAGATTGTGGTCGCAGTGTT | |  |
|  | 111028209 | uncharacterized protein | F: CCAGCATATGGCAAAGTCAA | | 183 |
|  |  |  | R: GGATGGACTGCTTGTTGGAT | |  |
|  | 111031319 | putative nuclease HARBI1 | F: TTAAAGGCTCGGTTCAGGTG | | 209 |
|  |  |  | R: CTTCCTCGAATGCCATCTTC | |  |
|  | 111035004 | AGLY_00338 | F: AAGGGCCAGCTTATCTCCAT | | 190 |
|  |  |  | R: GCCGTTTCATGTGGATTTTC | |  |
|  | 111036236 | troponin C-like | F: ACAACAACATGTCCGACGAA | | 168 |
|  |  |  | R: TTCAAAAAGTGCTGCGTTTG | |  |
|  | 111038117 | zinc finger MYM-type protein 1-like | F: TCATGAGCATTGAGCACCTC | | 215 |
|  |  |  | R: TAGCCGCACCACTAAGGACT | |  |
|  | 111037485 | uncharacterized protein | F: GCCAATCGCAAAAAGCTTAG | | 242 |
|  |  |  | R: AGCTGGCCCAGTTCAACTTA | |  |
|  | 111032528 | histone H3.2 | F: ACAGACCTGCGTTTCCAAAG | | 165 |
|  |  |  | R: ACGTTCTCCACGGATACGAC | |  |
|  | 111035341 | extensin-like isoform X1 | F: TACTCGTCACCGCACTTCAG | | 246 |
|  |  |  | R: CGATCGGTGGCGTTAGTAAT | |  |
|  | 100161709 | venom serine carboxypeptidase-like | F: AAACGGAAAGCACAAAATCG | | 225 |
|  |  |  | R: GGTTTTTCCACGAACTGCAT | |  |
|  | 111033337 | zinc finger and BTB domain-containing protein 24-like | F: TCCGTTGAAGGGTAAAGTCG | | 200 |
|  |  |  | R: AAGGTTGTCTCTCCGTGTGAA | |  |
|  | 111027646 | Actin binding protein 1 | F: AATTGGACGCGAACAAAAAC | | 153 |
|  |  |  | R: TTTCTTCGATTCCCCCTTCT | |  |
|  | 111032238 | 40S ribosomal protein S18 (PRS18) | F: CCTGTCCACCTTGTAGCGTT | | 132 |
|  |  |  | R: TGTGCTGGAACTTCTCAGGG  TGTGCTGGAACTTCTCAGGG  TGTGCTGGAACTTCTCAGGG | |  |
| RNAi | 111036118 | MpGABR | F: taatacgactcactatagggCTTGACGACGGGCAACTATT | | 325 |
|  |  |  | R: taatacgactcactatagggAATAGTTGCCCGTCGTCAAG | |  |
|  | 111041856 | MpGABRAP | F: taatacgactcactatagggTGAAGTTCCAGTACAAAG | | 183 |
|  |  |  | R: taatacgactcactatagggCTTTTTCTTGTCGAGCTCGC | |  |
|  | 111036117 | MpGABRB | F: taatacgactcactatagggCGAATTCATCCGGATACACC | | 250 |
|  |  |  | R: taatacgactcactatagggAGGGACACTTCGTTGGACAC | |  |
|  | ACY56286 | GFP | F: taatacgactcactatagggTCGGCGGCAATCCTGATCAA | | 476 |
|  |  |  | R: taatacgactcactatagggTCACAGGGTAAAATTCAGCA | |  |
| cloning | 111036118 | MpGABR | F: ATGACGTGTGGCGGCCGGCGG | | 2082 |
|  |  |  | R: TCAGTCTGCGCCCAGCAGCAC | |  |
|  | 111041856 | MpGABRAP | F: ATGAAGTTCCAGTACAAAGAA | | 354 |
|  |  |  | R: TCACACTCGTCCATAGACATTTTC | |  |
|  | 111036117 | MpGABRB | F: ATGACCGGCCGCGCCGCGCAC | | 750 |
|  |  |  | R: TCATACTTGCGTTGGAGATATTTT | |  |
| heterologous expression | 111036118 | MpGABR | F: CGCGGATCCGCGATGACGTGTGGCGGCCGGCGG | | 2105 |
|  |  |  | R: TCCCCCGGGGGTCAGTCTGCGCCCAGCAGCAC | |  |

| **Table S3.** Oligonucleotide sequences for PCR, the sgRNA synthesis template and the fragment of donor DNA for homology-directed repair (HDR) | |
| --- | --- |
| **Sequence name** | **Sequences (5’ to 3’)** |
| sgRNA-F (R122T) | TAATACGACTCACTATATTAGCGTATAGAAAACGACCT |
| sgRNA-R (R122T) | TTCTAGCTCTAAAACAGGTCGTTTTCTATACGCTAA |
| donor DNA (R122T) | GGACTTCACATTGGATTTTTACTTTCGTCAATTTTGGACCGATCCTCGTTTAGCGTATACAAAACGACCTGGTGTAGAAACACTATCGGTTGGATCAGAGTTCATTAAGAATATTTGGGTACCTGACACCTTTTTTGTAAATGAAAAACAATCATATTTTCACATTGCAACAACCAGTAATGAATTCATACGTGTGCATCATTCTGGATCGATAACAAGAAGTATTAG |
| Check-F | TACGTAAGTTCTCACTGCCAA |
| Check-R | GTTTGGATGTAGGGTTAGTTG |
| Note: The GenBank number of target sequence is KF881792.1 (*Drosophila melanogaster*). | |

**Table S4.** Screening of genes significantly differentially expressed in *M. persicae* in response to betulin at 48 h post treatment

| **Gene ID or GenBank No.** | **FPKM** | | **Log_2_ (fold change)** | **Description** |
| --- | --- | --- | --- | --- |
|  | **CK** | **betulin** |  |  |
| 111028237 | 0.03 | 6.54 | 7.62 | maltase 2-like |
| 111029701 | 0.09 | 4.05 | 5.44 | E3 SUMO-protein ligase KIAA1586-like isoform X1 |
| 111028209 | 0.03 | 0.97 | 5.03 | uncharacterized protein |
| 111028937 | 0.01 | 0.25 | 4.76 | uncharacterized protein |
| 111031319 | 0.04 | 1.15 | 4.73 | putative nuclease HARBI1 |
| 111032615 | 0.01 | 0.15 | 4.49 | AGLY_013951 |
| 111026557 | 0.01 | 0.24 | 4.19 | uncharacterized protein |
| 111035004 | 0.12 | 1.92 | 4.04 | AGLY_00338 |
| 111028447 | 3.86 | 62.79 | 4.02 | histone H4-like |
| 111036236 | 1.82 | 28.98 | 3.99 | troponin C-like |
| 111042749 | 0.11 | 1.54 | 3.81 | uncharacterized protein |
| 111030901 | 0.03 | 0.45 | 3.75 | protein ALP1-like |
| 111028166 | 0.66 | 7.98 | 3.59 | uncharacterized protein |
| 111042326 | 0.08 | 0.89 | 3.48 | uncharacterized protein |
| 111039160 | 0.04 | 0.41 | 3.36 | UDP-glucuronosyltransferase 2C1-like |
| 111032939 | 0.93 | 9.21 | 3.31 | uncharacterized protein |
| 111035373 | 1.67 | 16.50 | 3.30 | uncharacterized protein |
| 111028448 | 19.90 | 174.00 | 3.13 | histone H4 |
| 111038117 | 0.11 | 0.85 | 2.95 | zinc finger MYM-type protein 1-like |
| 111030763 | 0.10 | 0.79 | 2.93 | metalloendopeptidase OMA1, mitochondrial-like |
| 111035272 | 44.76 | 334.32 | 2.90 | cuticle protein 65 |
| 111042454 | 0.17 | 1.20 | 2.85 | uncharacterized protein |
| 111026977 | 0.07 | 0.52 | 2.82 | glucose dehydrogenase [FAD, quinone]-like |
| 111039754 | 16.06 | 109.52 | 2.77 | uncharacterized protein |
| 111042340 | 0.36 | 2.36 | 2.71 | 52 kDa repressor of the inhibitor of the protein kinase-like |
| 111038926 | 2876.35 | 18514.22 | 2.69 | uncharacterized protein |
| 111026270 | 1019.78 | 6215.78 | 2.61 | uncharacterized protein |
| 111032012 | 0.52 | 3.13 | 2.60 | uncharacterized protein |
| 111026880 | 0.55 | 3.20 | 2.53 | uncharacterized protein |
| 111036448 | 4.73 | 25.21 | 2.41 | flightin |
| 111030965 | 0.42 | 2.23 | 2.41 | uncharacterized protein |
| 111037485 | 0.14 | 0.72 | 2.33 | uncharacterized protein |
| 111028444 | 13.35 | 66.76 | 2.32 | histone H3.2 |
| 111042550 | 40.17 | 192.79 | 2.26 | BCL2/adenovirus E1B 19 kDa protein-interacting protein 3 |
| 111032531 | 3.02 | 14.47 | 2.26 | histone H4-like |
| 111037248 | 0.76 | 3.49 | 2.21 | uncharacterized protein |
| 111039704 | 0.24 | 1.09 | 2.20 | zinc finger BED domain-containing protein 1-like |
| 111028445 | 8.90 | 40.04 | 2.17 | histone H2B-like |
| 27109199 | 57.14 | 237.99 | 2.06 | cytochrome c oxidase subunit 3 |
| 111032559 | 2.83 | 11.69 | 2.05 | histone H2B-like |
| 111029588 | 0.66 | 2.74 | 2.04 | uncharacterized protein |
| 111027021 | 4.90 | 19.79 | 2.01 | AGLY_017505 |
| 111032546 | 7.90 | 30.86 | 1.97 | FQR65_LT12436 |
| 111040339 | 7.81 | 27.66 | 1.82 | AGLY_009050 |
| 111039904 | 0.61 | 2.13 | 1.81 | uncharacterized protein |
| 111032555 | 5.81 | 20.12 | 1.79 | histone H2A |
| 111028488 | 0.28 | 0.92 | 1.74 | enkurin-like |
| 111032107 | 0.11 | 0.37 | 1.72 | piggyBac transposable element-derived protein 3-like |
| 111032545 | 6.74 | 22.19 | 1.72 | histone H2A-like |
| 111031834 | 0.37 | 1.20 | 1.71 | uncharacterized protein |
| 111032543 | 4.45 | 14.50 | 1.70 | histone H3.2 |
| 111039139 | 1.04 | 3.34 | 1.68 | AGLY_010144 |
| 111035978 | 0.59 | 1.88 | 1.67 | uncharacterized protein |
| 111031173 | 13.97 | 44.09 | 1.66 | uncharacterized protein |
| 111029243 | 51.80 | 162.59 | 1.65 | uncharacterized protein |
| 111038156 | 1.97 | 6.18 | 1.65 | membrane-associated guanylate kinase |
| 111032541 | 0.45 | 1.39 | 1.63 | histone H1B, sperm-like |
| 111035832 | 1.12 | 3.48 | 1.63 | nose resistant to fluoxetine protein 6-like |
| 111042476 | 6.46 | 19.78 | 1.61 | histone H1A, sperm-like |
| 111027761 | 5.60 | 16.96 | 1.60 | cathepsin B-like |
| 111028442 | 6.26 | 18.76 | 1.58 | histone H1A, sperm-like |
| 111033017 | 14.77 | 43.33 | 1.55 | uncharacterized protein |
| 111030609 | 2.60 | 7.52 | 1.53 | uncharacterized protein |
| 111032539 | 5.91 | 16.93 | 1.52 | histone H1A, sperm-like |
| 111028446 | 47.73 | 136.54 | 1.52 | histone H2A-like isoform X1 |
| 111042602 | 18.47 | 52.80 | 1.52 | pancreatic triacylglycerol lipase |
| 111029483 | 5.12 | 14.63 | 1.52 | histone H2A |
| 111036169 | 0.47 | 1.33 | 1.50 | AGLY_013443 |
| 111032528 | 10.97 | 30.80 | 1.49 | histone H3.2 |
| 111030632 | 291.86 | 812.42 | 1.48 | regucalcin-like isoform X1 |
| 111042439 | 0.50 | 1.38 | 1.47 | adenylate kinase 7-like |
| 111042522 | 2.63 | 7.26 | 1.47 | uncharacterized protein |
| 111031729 | 14.12 | 38.19 | 1.44 | histone H2B |
| 111038355 | 0.18 | 0.48 | 1.43 | pseudouridine-5'-phosphatase-like |
| 111032561 | 8.07 | 21.69 | 1.43 | histone H2A-like |
| 111031569 | 55.31 | 146.43 | 1.40 | dnaJ homolog subfamily B member 13-like |
| 111034497 | 2.09 | 5.47 | 1.39 | agglutinin-like protein 3 isoform X1 |
| 111033668 | 115.94 | 302.99 | 1.39 | asparagine synthetase |
| 111027227 | 0.47 | 1.23 | 1.38 | THAP domain-containing protein 1-like isoform X |
| 111030094 | 2.01 | 5.25 | 1.38 | uncharacterized protein |
| 111030036 | 1.29 | 3.36 | 1.38 | probable cytochrome P450 6a13 isoform X1 |
| 111039664 | 0.28 | 0.74 | 1.38 | inter-alpha-trypsin inhibitor heavy chain H3 isoform X2 |
| 111041393 | 0.54 | 1.39 | 1.37 | uncharacterized protein |
| 111027567 | 0.87 | 2.24 | 1.37 | AGLY_016272 |
| 111041770 | 2.31 | 5.95 | 1.36 | putative nuclease HARBI1 |
| 111028079 | 2.48 | 6.35 | 1.35 | uncharacterized protein |
| 111042514 | 0.48 | 1.22 | 1.35 | UDP-glucuronosyltransferase 2B2-like |
| 111027018 | 0.59 | 1.50 | 1.34 | uncharacterized protein |
| 111031730 | 13.23 | 33.48 | 1.34 | histone H2A-like |
| 111036414 | 21.59 | 54.17 | 1.33 | cathepsin B-like |
| 111042600 | 13.60 | 34.06 | 1.32 | prostatic acid phosphatase-like isoform X2 |
| 111029718 | 15.36 | 38.35 | 1.32 | UDP-glucuronosyl transferase 344E7 |
| 111042513 | 0.67 | 1.66 | 1.31 | BAC VMRC38-20-A9 |
| 111027070 | 0.46 | 1.13 | 1.31 | uncharacterized protein |
| 111033939 | 10.22 | 24.93 | 1.29 | uncharacterized protein |
| 111037298 | 2.88 | 6.87 | 1.25 | uncharacterized protein |
| 111036411 | 24.06 | 56.60 | 1.23 | cathepsin B-like |
| 111033556 | 18.99 | 44.62 | 1.23 | cytochrome P450 6k1 |
| 111031644 | 13.72 | 32.18 | 1.23 | fatty acyl-CoA reductase wat-like isoform X1 |
| 111030182 | 2.39 | 5.60 | 1.23 | uncharacterized protein |
| 111034608 | 4.06 | 9.43 | 1.21 | uncharacterized protein |
| 27109224 | 12.16 | 28.20 | 1.21 | NADH dehydrogenase subunit 4 |
| 111029998 | 2.52 | 5.83 | 1.21 | uncharacterized protein |
| 111033168 | 35.53 | 82.01 | 1.21 | AGLY_004239 |
| 111040065 | 0.33 | 0.76 | 1.19 | BLOC-1-related complex subunit 8 homolog |
| 111039739 | 14.12 | 32.17 | 1.19 | ANTAGONIST OF LIKE HETEROCHROMATIN PROTEIN 1-like |
| 111031862 | 2.84 | 6.41 | 1.18 | putative malate dehydrogenase 1B |
| 111040667 | 146.48 | 327.35 | 1.16 | fatty acyl-CoA reductase wat-like |
| 111032544 | 10.21 | 22.81 | 1.16 | histone H2B-like |
| 111039710 | 62.64 | 138.34 | 1.14 | uncharacterized protein |
| 111036825 | 0.73 | 1.60 | 1.14 | hypothetical protein AGLY_009353 |
| 111030990 | 1.05 | 2.31 | 1.13 | zinc finger BED domain-containing protein 1-like isoform X1 |
| BK010800.1 | 0.44 | 0.96 | 1.13 | cytochrome P450 monooxygenase CYP6CY49 |
| 111030846 | 1.83 | 3.89 | 1.09 | uncharacterized protein |
| 111035602 | 0.51 | 1.08 | 1.07 | nose resistant to fluoxetine protein 6-like isoform X1 |
| 103309211 | 1.00 | 2.10 | 1.07 | putative nuclease HARBI1 |
| 27109198 | 39.78 | 83.37 | 1.07 | ATP synthase F0 subunit 6 |
| 111043077 | 3.08 | 6.40 | 1.06 | uncharacterized protein |
| 111032913 | 2.56 | 5.31 | 1.05 | cathepsin B-like |
| 111031087 | 3.76 | 7.80 | 1.05 | ANTAGONIST OF LIKE HETEROCHROMATIN PROTEIN 1-like |
| KAF0749006.1 | 0.62 | 1.29 | 1.05 | uncharacterized protein |
| 107168481 | 3.61 | 7.47 | 1.05 | uncharacterized protein |
| 111040079 | 1.79 | 3.68 | 1.04 | ADP-ribosylation factor-like protein 13B isoform X1 |
| 111040738 | 0.73 | 1.51 | 1.04 | zinc finger MYM-type protein 1-like |
| 111036924 | 1.46 | 2.98 | 1.03 | uncharacterized protein |
| 111042142 | 0.83 | 1.70 | 1.03 | ring-infected erythrocyte surface antigen-like |
| 111026458 | 2.67 | 5.41 | 1.02 | Integrase catalytic domain-containing protein |
| 111028300 | 1.21 | 2.44 | 1.02 | Transposable element P transposase |
| 111036037 | 15030.14 | 30585.48 | 1.02 | uncharacterized protein |
| 111029334 | 4.92 | 10.12 | 1.04 | uncharacterized protein |
| 111027522 | 82.17 | 40.95 | -1.00 | alpha-tocopherol transfer protein-lik |
| 111036239 | 125.08 | 62.34 | -1.00 | aspartate aminotransferase, cytoplasmic-like isoform X1 |
| 111027535 | 5.52 | 2.73 | -1.02 | plexin-A3 |
| 111035341 | 24.67 | 12.12 | -1.02 | extensin-like isoform X1 |
| 111030420 | 46.53 | 22.80 | -1.03 | nose resistant to fluoxetine protein 6 |
| 111037974 | 222.38 | 108.77 | -1.03 | uncharacterized protein |
| 111038225 | 5.68 | 2.77 | -1.04 | uncharacterized protein |
| 111039266 | 223.21 | 108.18 | -1.04 | dopamine N-acetyltransferase-like |
| 111042475 | 4.98 | 2.39 | -1.06 | nose resistant to fluoxetine protein 6 |
| 111034181 | 0.67 | 0.32 | -1.06 | uncharacterized protein |
| 111029593 | 34.64 | 16.35 | -1.08 | thiamine transporter 2-like |
| 111034899 | 21.55 | 10.08 | -1.10 | aminopeptidase N 2 |
| 111030691 | 12.76 | 5.84 | -1.13 | protein slit-like |
| 111029021 | 323.56 | 147.78 | -1.13 | omega-amidase NIT2-like |
| 111030483 | 1.03 | 0.47 | -1.14 | esterase E4-like |
| 111031195 | 9.99 | 4.49 | -1.15 | cytochrome P450 4C1-like |
| 111036117 | 779.66 | 345.76 | -1.17 | MpGABRB |
| 111033226 | 6.73 | 2.97 | -1.18 | facilitated trehalose transporter Tret1-like |
| 111030384 | 267.79 | 118.14 | -1.18 | uncharacterized protein |
| 111031854 | 96.18 | 41.09 | -1.23 | kynurenine/alpha-aminoadipate aminotransferase, mitochondrial-like |
| 111035008 | 4.86 | 1.99 | -1.29 | AGLY_014733 |
| 111028477 | 4.18 | 1.70 | -1.30 | 2-hydroxyacylsphingosine 1-beta-galactosyltransferase-like |
| 111041856 | 3.73 | 1.51 | -1.30 | MpGABRAP |
| 111029126 | 87.43 | 34.49 | -1.34 | alpha-tocopherol transfer protein-like |
| 111033229 | 3.37 | 1.26 | -1.42 | facilitated trehalose transporter Tret1-like isoform X1 |
| 111042433 | 2151.40 | 782.70 | -1.46 | heat shock protein 70 A1-like |
| 111028805 | 41.49 | 14.51 | -1.52 | high affinity cationic amino acid transporter 1-like |
| 100161709 | 7.20 | 2.44 | -1.56 | venom serine carboxypeptidase-like |
| 111040524 | 12.63 | 4.23 | -1.58 | cytochrome P450 4C1-like |
| 111032647 | 6.42 | 2.13 | -1.59 | uncharacterized protein |
| 111041470 | 1.42 | 0.46 | -1.63 | zinc finger BED domain-containing protein 1-like |
| 111031401 | 16.98 | 4.98 | -1.77 | protein 5NUC isoform X2 |
| 111040367 | 0.76 | 0.20 | -1.91 | glucose dehydrogenase [FAD, quinone] isoform X1 |
| 111033337 | 4.46 | 1.01 | -2.14 | zinc finger and BTB domain-containing protein 24-like |
| 111035764 | 0.66 | 0.15 | -2.18 | bifunctional lycopene cyclase/phytoene synthase |
| 111031197 | 3.18 | 0.58 | -2.45 | cytochrome P450 CYP380C7 |
| 111036240 | 4.64 | 0.78 | -2.57 | aspartate aminotransferase, cytoplasmic-like |
| 111034158 | 0.36 | 0.06 | -2.58 | uncharacterized protein |
| 111028612 | 0.31 | 0.05 | -2.72 | scoloptoxin SSD14-like |
| 111036118 | 0.66 | 0.05 | -3.83 | MpGABR |
| 111027646 | 2.15 | 0.07 | -4.94 | Actin binding protein 1 |

**Table S5.** Complete sequence information for the *GABA_A_ receptor* gene of *M. persicae*

| **Gene** | **Gene ID** | **Coding sequence (bp)** | **Deduced amino acid** | **Molecular weight (kDa)** | **Isoelectric point** |
| --- | --- | --- | --- | --- | --- |
| MpGABR | 111036118 | 2082 | 693 | 77.20 | 9.85 |
| MpGABRAP | 111041856 | 357 | 118 | 14.08 | 9.63 |
| MpGABRB | 111036117 | 753 | 250 | 28.01 | 7.20 |

**Table S6.** Sequences and relevant information for phylogenetic analysis of GABA_A_ receptor

| **Species** | **Accession or GenBank No.** | **Order** |
| --- | --- | --- |
| *Myzus persicae* | XP_022173711.1 | Hemiptera |
| *Metopolophium dirhodum* | XP_060864885.1 | Hemiptera |
| *Acyrthosiphon pisum* | XP_008183008.2 | Hemiptera |
| *Macrosiphum euphorbiae* | CAI6365831.1 | Hemiptera |
| *Rhopalosiphum padi* | XP_060842789.1 | Hemiptera |
| *Melanaphis sacchari* | XP_025199515.1 | Hemiptera |
| *Aphis gossypii* | XP_050054462.1 | Hemiptera |
| *Diuraphis noxia* | XP_015376355.1 | Hemiptera |
| *Sipha flava* | XP_025404819.1 | Hemiptera |
| *Cinara cedri* | VVC42195.1 | Hemiptera |
| *Adelges cooleyi* | XP_050429873.1 | Hemiptera |
| *Laodelphax striatellus* | RZF38914.1 | Hemiptera |
| *Macrosteles quadrilineatus* | XP_054276917.1 | Hemiptera |
| *Planococcus citri* | XP_065213924.1 | Hemiptera |
| *Cyrtorhinus lividipennis* | AHW29556.1 | Hemiptera |
| *Bemisia tabaci* | XP_018901515.2 | Hemiptera |
| *Cimex lectularius* | XP_024083544.1 | Hemiptera |
| *Halyomorpha halys* | XP_014281964.1 | Hemiptera |
| *Anopheles gambiae* | XP_001688775.1 | Diptera |
| *Aedes aegypti* | XP_021696320.1 | Diptera |
| *Sitodiplosis mosellana* | XP_055320038.1 | Diptera |
| *Culicoides brevitarsis* | XP_063702909.1 | Diptera |
| *Lutzomyia longipalpis* | XP_055686441.1 | Diptera |
| *Ceratitis capitata* | XP_004525611.1 | Diptera |
| *Phlebotomus papatasi* | XP_055701411.1 | Diptera |
| *Drosophila persimilis* | XP_026849499.1 | Diptera |
| *Drosophila melanogaster* | AHE41089.1 | Diptera |
| *Drosophila simulans* | AAK00512.1 | Diptera |
| *Plutella xylostella* | NP_001292463.1 | Lepidoptera |
| *Papilio xuthus* | XP_013179056.1 | Lepidoptera |
| *Papilio polytes* | XP_013133342.1 | Lepidoptera |
| *Pieris rapae* | XP_022123609.1 | Lepidoptera |
| *Danaus plexippus* | XP_032527541.1 | Lepidoptera |
| *Amyelois transitella* | XP_013189102.1 | Lepidoptera |
| *Ostrinia furnacalis* | XP_028160229.1 | Lepidoptera |
| *Chilo suppressalis* | ASY91958.1 | Lepidoptera |
| *Trichoplusia ni* | XP_026735745.1 | Lepidoptera |
| *Heliothis virescens* | AAB62572.1 | Lepidoptera |
| *Spodoptera frugiperda* | XP_035445711.1 | Lepidoptera |
| *Bombyx mandarina* | XP_028032124.1 | Lepidoptera |
| *Bombyx mori* | NP_001093294.1 | Lepidoptera |
| *Manduca sexta* | XP_030028507.1 | Lepidoptera |
| *Frankliniella occidentalis* | XP_026285852.1 | Thysanoptera |
| *Thrips palmi* | XP_034255608.1 | Thysanoptera |
| *Neodiprion lecontei* | XP_015509151.1 | Hymenoptera |
| *Athalia rosae* | XP_012264889.1 | Hymenoptera |
| *Cephus cinctus* | XP_015589073.1 | Hymenoptera |
| *Orussus abietinus* | XP_023289674.1 | Hymenoptera |
| *Chelonus insularis* | XP_034938828.1 | Hymenoptera |
| *Microplitis demolitor* | XP_008549804.1 | Hymenoptera |
| *Nasonia vitripennis* | XP_016845786.2 | Hymenoptera |
| *Ceratosolen solmsi marchali* | XP_011495770.1 | Hymenoptera |
| *Camponotus floridanus* | XP_011252265.1 | Hymenoptera |
| *Harpegnathos saltator* | XP_011137565.1 | Hymenoptera |
| *Megachile rotundata* | XP_012143665.1 | Hymenoptera |
| *Osmia lignaria* | XP_034181602.1 | Hymenoptera |
| *Eufriesea mexicana* | XP_017755009.1 | Hymenoptera |
| *Apis dorsata* | XP_006619002.1 | Hymenoptera |
| *Apis mellifera* | XP_006565169.1 | Hymenoptera |
| *Bombus impatiens* | XP_024220796.1 | Hymenoptera |
| *Agrilus planipennis* | XP_018318732.1 | Coleoptera |
| *Photinus pyralis* | XP_031346787.1 | Coleoptera |
| *Nicrophorus vespilloides* | XP_017787270.1 | Coleoptera |
| *Onthophagus taurus* | XP_022907694.1 | Coleoptera |
| *Tribolium castaneum* | NP_001107809.1 | Coleoptera |
| *Tribolium madens* | XP_044271991.1 | Coleoptera |
| *Aethina tumida* | XP_019870942.1 | Coleoptera |
| *Anoplophora glabripennis* | XP_018564077.1 | Coleoptera |
| *Leptinotarsa decemlineata* | XP_023012203.1 | Coleoptera |
| *Oulema oryzae* | BAM66322.1 | Coleoptera |

**Table S7.** Binding energy and nonbonding interactions between betulin and GABA_A_ receptor

| **Compound** | **Binding Energy (**kcal⋅mol^–1^**)** | **van der Waals** | **H-Bond** | | **Hydrophobic Interaction** | |
| --- | --- | --- | --- | --- | --- | --- |
|  |  |  | **Amino Acid-**  **Ligand Atom** | **Distance (Å)** | **Amino Acid-Ligand Atom** | **Distance (Å)** |
| betulin | -6.38 | PHE172, GLY174, TYR176 | ALA226 [C…H]  THR228 [C-O.H] | 3.31  2.16 | ARG224 [Alkyl]  ALA226 [Alkyl]  ALA226 [Alkyl]  PHE227 [Pi–Alkyl]  PHE227 [Pi–Alkyl] | 4.29  4.36  5.11  4.16  5.32 |

**Table S8.** Parameters of dose-response curves from microscale thermophoresis experiments.

| **Compound** | | **Strains** | ***K_d_* (****μM)** | **95% CI ^a^ (μM)** | **HillSlope** | ***r^2^*** |
| --- | --- | --- | --- | --- | --- | --- |
|  | betulin | WT | 2.24 | 1.860-2.687 | 0.0726 | 0.9864 |
|  |  | R224A | 2.31 | 1.799-2.753 | 0.0827 | 0.9822 |
|  |  | A226T | 2.27 | 1.958-2.628 | 0.0538 | 0.9918 |
|  |  | F227Y | 2.25 | 1.860-2.690 | 0.0823 | 0.9850 |
|  |  | T228R | 5321 | 4599-6152 | 0.0560 | 0.9916 |

CI ^a^, Confidence Interval; *R^2^*, coefficient of determination.

**Table S9.** LD_50_ values of betulin and pymetrozine against *D. melanogaster* at 72 h

| **Compounds** | **Strains** | **Regression**  **equation** | **LD_50_**  **(μg⋅fly^−1^)** | **95% Confidence**  **Interval (μg⋅fly^−1^)** | ***r^2^*** |
| --- | --- | --- | --- | --- | --- |
| betulin | WT | - | > 1000 ^a^ | - | - |
|  | R122T | Y=3.6687+0.9282X | 27.1853 | 25.8872-28.5485 | 0.9944 |
| pymetrozine | WT | Y=5.8681+1.3720X | 0.2330 | 0.2311-0.2348 | 0.9933 |
|  | R122T | Y=5.9388+1.4937X | 0.2352 | 0.2335-0.2369 | 0.9903 |

^a^ Toxicity bioassays were not performed with concentrations exceeding 1000 μg⋅fly^−1^.
